# Context and Identity Cues Mediate Individual Recognition in a Highly Social Cichlid Fish

**DOI:** 10.64898/2026.09.27.754830

**Authors:** Emily K. Lessig, Isha Kalpaguri, Abiel Issac, Hans A. Hofmann

**Author notes:** Dr. Emily Lessig GIMM- Gulbenkian Institute for Molecular Medicine, Rua Quinta Grande 6, 2780-156 Oeiras, Portugal.

## Abstract

Social animals must identify, evaluate, and appropriately respond to other individuals, often relying on multiple sources of information to guide context-dependent behavior. While individual recognition is thought to be widespread and critical for many social interactions (e.g. territoriality, dominance hierarchies), convincing demonstrations across animals beyond class-level recognition remain rare. Further, the cues animals use to recognize individuals remains poorly understood, especially in natural environments where they are exposed to numerous cues which may require cue integration. Here, we examine how socially dominant males of the African cichlid fish, *Astatotilapia burtoni,* use individual recognition to modulate aggression and whether this process depends on the spatial context. Using naturalistic experimental setups, we exposed males to familiar and unfamiliar neighbors while systematically manipulating spatial location and presenting a neighbor either singly or simultaneously to simulate competing social cues. Additionally, we measured circulating testosterone to assess endocrine correlates of recognition and aggression. We found that focal males habituated to familiar neighbors only when they remained in a consistent territorial location, demonstrating that spatial context is important for recognition. Further, when familiar and unfamiliar neighbors were presented simultaneously, males preferentially aggressed toward unfamiliar rival and adjust their aggressive response to the perceived threat of their neighbors, indicating that identity-based threat assessment is maintained under competing social demands. Together, our results show that recognition is a flexible, multi-faceted process that depends on the integration of identity and contextual information to guide territorial decision-making under dynamic social conditions.

## INTRODUCTION

Animals that live in social groups often need to recognize other individuals – such as kin, mates, neighbors, rivals, other conspecifics, or predators – to engage in context-appropriate behavior in almost all their social interactions. Even though these kinds of recognition can rely on incidental cues that have evolved independent of any recognition context, individuals in some species may also signal their identity when it is beneficial to be recognized. For example, daffodil cichlids (*Neolamprologus pulcher*) and paper wasps (*Polistes fuscatus*) can distinguish individuals using facial color patterns or markings (Kohda et al., 2015; Tibbetts, 2002). Recognition in such cases involves a sender-receiver relationship that requires both the sender to broadcast a signal that is identifiable and the receiver to learn and remember the identity of the signaler (Johnstone 1997; Dale et al. 2001; Beecher 1989). Selection, therefore, acts on both signalers and receivers – signalers need to produce identifiable and memorable signals and receivers need to possess the cognitive capacity that facilitates recognition. Identity signals can have different properties (e.g. quality, variability, modality, but see Dale 2006; Tibbetts and Dale 2007) and can be received through a range of sensory modalities including visual, auditory, olfactory, or tactile cues, although likely in combination (Hebets & Papaj, 2005). Consequently, to avoid the costs of misidentifying individuals, many organisms have evolved sophisticated recognition mechanisms such as individual recognition, a special case of recognition that occurs when an organism identifies another according to its individually distinctive characteristics (Dale et al. 2001; Tibbetts and Dale 2007).

As selection can strongly favor recognition abilities, many species have evolved various sensory specializations to help them identify individuals more easily. Vocal recognition is found across a broad range of taxa (reviewed by Carlson et al. 2020), including frogs (Bee & Gerhardt, 2002; Chuang et al., 2017; Lesbarrères & Lodé, 2002), birds (Gentner & Hulse, 1998; Robertson, 1996; Wunderle, 1978), fish (Myrberg & Riggio, 1985), and a variety of mammals (Arnold & Wilkinson, 2011; Briseño-Jaramillo et al., 2015; Espmark, 1971). For example, emperor penguins (*Aptenodytes forsteri*) find their chicks among thousands of unrelated individuals after returning from long foraging trips using a two-voice system (Aubin et al., 2000). Olfactory cues play a large role in recognition (reviewed by Smadja and Butlin 2009; Wyatt 2003) with well-known examples in insects that use diverse pheromonal compounds for communication (Fleischer & Krieger, 2018) and rodents who use major urinary proteins to recognize individuals (McLennan, 2004).

Additionally, a variety of organisms use visual cues for recognition such as size, coloration, patterns, and markings. Facial recognition, for example, is common in humans (Tanaka & Farah, 1993), non-human primates (e.g. chimpanzees: Martin-Malivel and Okada 2007; rhesus monkeys: Rosenfeld and Van Hoesen 1979), sheep (Kendrick et al., 1996) and has even been described in wasps (Tibbetts, 2002, 2025). Some teleost fish species may have visual acuity comparable to humans (Douglas & Djamgoz, 2012) and can use facial coloration to distinguish between familiar and unfamiliar individuals, such as the damselfish, *Pomacentrus amboinensis* (Siebeck et al. 2010), the daffodil cichlid, *N. pulcher* (Kohda et al. 2015), and the blue discus cichlid, *Symphysopdon aequifasciatus* (Satoh et al. 2016). Facial recognition, a special case of individual recognition, can be advantageous when there are repeated interactions between the same individuals, especially in smaller sized groups, and can help mediate territorial aggression, moderate aggressive competition, and stabilize social dominance hierarchies (Tibbetts & Dale, 2007).

Some animals minimize the costs of conflict by using signals of fighting ability to directly assess and avoid rivals (e.g. large weapons; Guilford & Dawkins, 1995; Hofmann & Schildberger, 2001; Tibbetts et al., 2022). An individual’s fighting ability can also be assessed indirectly by observing pairwise interactions and remembering the outcome of those interactions such as through eavesdropping or transitive inference (Bonnie & Earley, 2007; Peake, 2005; Valone, 2007; Vasconcelos, 2008). For example, males of the Burton’s Mouthbrooder cichlid fish (*Astatotilapia burtoni*) can successfully make inferences on a social dominance hierarchy by watching pairwise fights between rival males (Grosenick et al. 2007). These studies suggest identifiable individual features (i.e., “identity cues”) are very important to assess an organism’s dominance status or competitive ability and may be used instead of engaging in direct competition.

Individual recognition can also reduce the costs of territorial defense by enabling territory holders to focus their aggressive efforts on non-territorial intruders rather than familiar neighbors (‘dear enemy effect’: Ydenberg et al. 1988; Temeles 1994). This ‘dear enemy’ effect has been demonstrated across a wide variety of species (reviewed by Werba et al. 2022) with well-known examples in territorial songbirds (Godard 1991; Akçay et al. 2009; Olendorf et al. 2004), fiddler crabs, *Uca mjoebergi*, (Backwell and Jennions 2004) and *A. burtoni* (Weitekamp and Hofmann 2017). In some species, however, territorial males increase aggression towards familiar neighbors based on their neighbor’s unpredictable behavior, which can pose a threat and require increased vigilance (i.e. ‘nasty neighbors’: Olendorf et al. 2004; Müller and Manser 2007) such as in the banded mongoose (*Mungos mungo*), who aggress more towards familiar compared to unfamiliar neighbors to reduce the risk of being cuckolded (Müller & Manser, 2007). This increased aggression towards a familiar neighbor could be due to neighbors moving beyond their established boundary lines (e.g. in the ‘incorrect’ territory) where the threat-level hypothesis predicts that territory owners will adjust their aggression levels according to the threat level of their neighbors (Temeles 1994). Specifically, familiar neighbors repositioning their territories could be perceived as reneging on the ‘dear enemy’ agreement and therefore as a threat. For example, hooded warblers (*Wilsonia citrina*) ignore the playbacks of neighbors positioned at the ‘correct’ territorial boundary, but aggress to playbacks of neighbors positioned in the ‘incorrect’ territorial boundary and to stranger calls (Godard 1991). Daffodil cichlids (*N. pulcher*) also show limited aggression towards familiar neighbors, but increase aggression towards familiar neighbors in unfamiliar territories (Saeki et al., 2018). These studies suggest that the location of a neighbor can also strongly influence how they are being perceived, and animals may be using context cues – in addition to or instead of identity cues – to recognize and respond to neighboring males.

Androgens such as testosterone mediate aggressive behavior and are typically released in response to a social challenge (Wingfield et al. 1990; Goymann et al. 2019). Surges in testosterone related to male-male encounters can also reinforce learning associated with aggressive encounters (Marler et al. 2005) as well as have rewarding properties that can elicit a conditioned place preference for the location in which a given encounter occurred (Marler et al. 2005; Gleason et al. 2009).

Although previous studies have demonstrated dear enemy relationships and context-dependent territorial behavior across a variety of taxa (Godard, 1991; Saeki et al., 2018; Temeles, 1994; Werba et al., 2022), it remains unclear whether these responses are mediated by individual recognition or broader forms of familiarity-based discrimination (e.g. class-level recognition). Additionally, how animals integrate multiple recognition cues simultaneously when evaluating rivals remains poorly understood. For example, whether identity and spatial context interact to shape territorial aggression and how animals prioritize conflicting social information is of particular interest. Moreover, while androgens such as testosterone are known to mediate aggressive behavior during social challenges, whether endocrine responses differ according to familiarity, identity, or contextual recognition cues is less well known.

Burton’s Mouthbrooder cichlid, *Astatotilapia burtoni,* is a highly social teleost fish species characterized by distinct and reversible dominant and subordinate phenotypes that encompass patterns of behavior, physiology, and neurobiology. Dominant *A. burtoni* males exhibit high levels of aggression and territorial defense that allow them to gain reproductive opportunities with females as well as bright coloration and high levels of sex steroids (Maruska 2014; Maruska et al. 2013). In their natural habitat, *A. burtoni* males form dense, lek-like communities where dominant males aggressively defend territories (Fernald and Hirata 1977; Fernald 1977). It is likely, therefore, that males would have repeated interactions with the same individuals where recognition mechanisms would be important to reduce the costs of agonistic competition. Additionally*, A. burtoni* males display context-dependent behavior that can be robustly quantified and manipulated in naturalistic laboratory settings (Maruska and Fernald 2018). For example, males modify their behavior by observing other individuals (Grosenick et al. 2007; Desjardins et al. 2012) such as modifying aggression based on presence or absence (Desjardins et al. 2012), or the identity (familiar or unfamiliar: Weitekamp et al. 2017; Weitekamp and Hofmann 2017) of other males. However, it remains unclear whether these responses are mediated by individual recognition and whether *A. burtoni* males integrate multiple recognition cues during rival assessment. In particular, whether identity and spatial context interact to shape territorial aggression and how males prioritize conflicting social information has not been directly tested.

In the present study, we used *A. burtoni* males to test whether dominant males use individual recognition to distinguish familiar vs. unfamiliar neighbors and the recognition cues that are involved. Specifically, we exposed males to neighbors that were either familiar or unfamiliar and varied their location (e.g. territories) and measured their aggressive responses. We hypothesized that focal males would respond most aggressively to unfamiliar neighbors. Additionally, if males were using individual recognition, we predicted they would show elevated levels of aggression to a familiar neighbor in an ‘incorrect’ territory, with lowest levels of aggression to a familiar neighbor in the ‘correct’ territory. We also quantified circulating testosterone levels in response to neighbor aggression and predicted males would differ in their hormonal responses across treatments, with highest levels of testosterone in response to an unfamiliar neighbor.

## MATERIALS AND METHODS

### Animals

*Astatotilapia burtoni* descended from a wild-caught stock population were kept in 100 l aquaria under conditions mimicking their natural environment. Fish were fed daily with cichlid flakes (Omega One Cichlid Flake Fish Food, Arcata Pet Supplies, Arcata, CA) and a 12:12 h light: dark cycle was maintained. All work was approved by the Institutional Animal Care and Use Committee at the University of Texas at Austin.

### Behavior Trials

Experimental tanks (100 l) were divided into 3 even compartments using acrylic dividers with 1 dominant male housed in the middle compartment (focal male) with 2 juveniles and 2 females, and 1 dominant male housed in a side compartment (neighbor male) with 2 juveniles and 2 females. Males housed in the same tank therefore remained physically separated throughout the experiment but were given visual exposure to neighbors during the habituation and stimulus treatments. Males were taken from communities where they had been dominant for at least 1 week prior to the start of experiments and neighbors were randomly assigned to either side compartment. Males were given halved terra-cotta pots to serve as bowers and females and juveniles were included in tanks to elicit naturalistic male behavior. Males underwent a 7-day acclimation period where opaque acrylic dividers were placed between dominant males for visual isolation. Males then underwent a 4-day habituation period where they were given visual exposure to each other for 1 h twice a day (between 8:30–10:30h) through clear acrylic dividers (Fig. 1).

**Figure 1.**
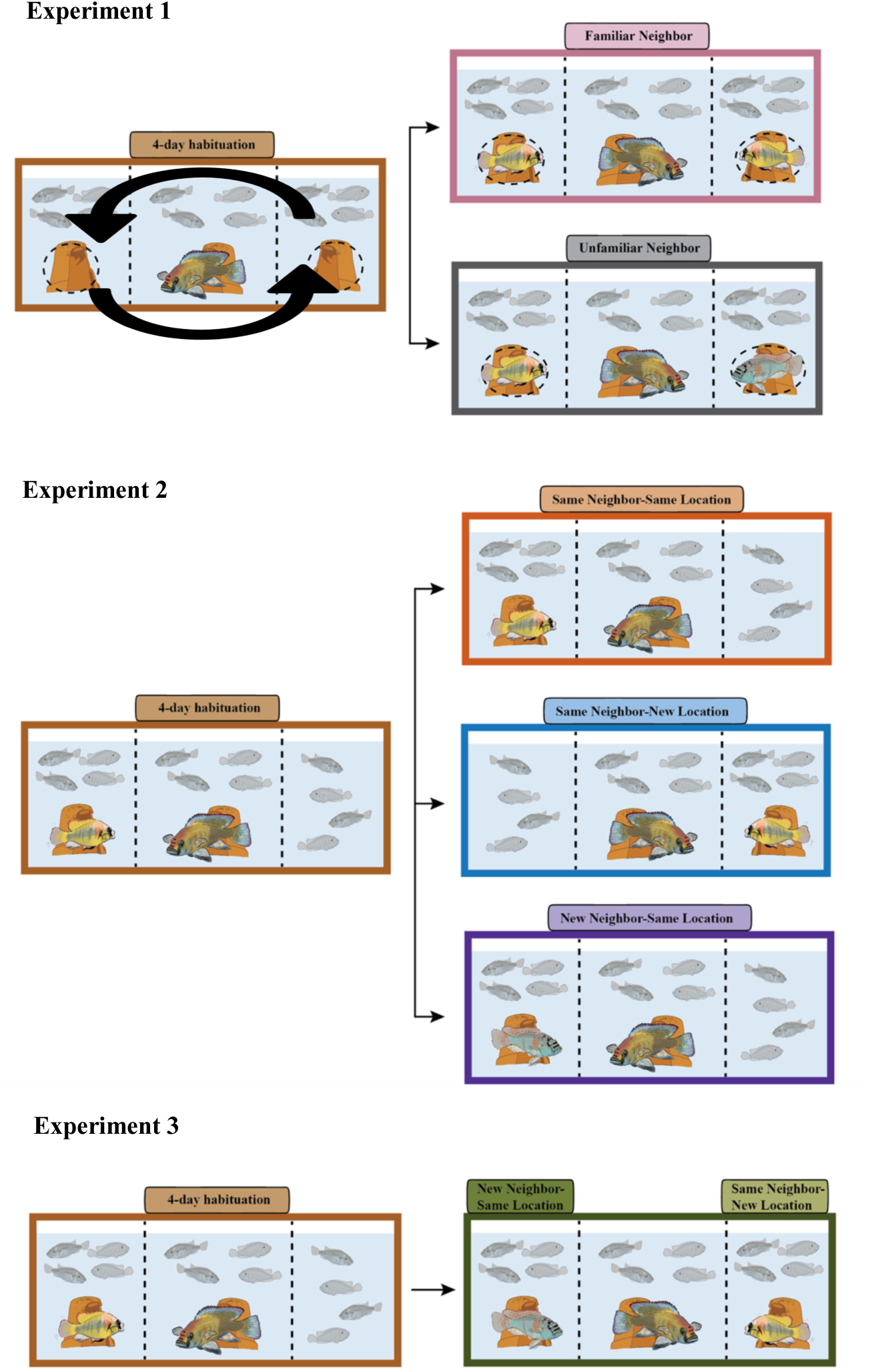
Experimental paradigm. Dominant *A. burtoni* males were given a 7-day acclimation period in experimental tanks with 2 juveniles and 2 females. Males then underwent a 4-day habituation period where they were given visual exposure to a neighboring dominant male for 1h twice a day. Experiment 1: Males experienced a 4-day habituation period where i) the same (familiar) neighbor was used across all 4 days and moved compartments daily ii) a new (unfamiliar) neighbor was used across all 4 days and moved compartments daily. Experiment 2: Following the habituation period, focal males were exposed to one of three single stimulus treatments that varied in the identity of their neighbor (familiar vs. unfamiliar) and/or their location (same vs. different location) or were presented with two neighbors simultaneously (Experiment 3). Fish images by C.N. Friesen.

In Experiment 1, the neighboring male and/or his location in the tank varied across the 4-day habituation period. In Experiment 2 and 3, the neighboring male was the same individual, in the same location across the 4-day habituation period. For treatments requiring males to be moved to a new location, animals were transferred using a dip net and given an overnight acclimation period prior to experimental testing. Additionally, all males were size-matched. The standard length (mm) and body mass (g) were measured, the gonadosomatic index (GSI) was calculated ((gonad weight (g) / body mass (g)) × 100), and body color (classified as blue, yellow or blue-yellow) was assessed for each dominant male. Standard lengths ranged from 45–60 mm and body mass from 1.80–5.37 g, with GSI ranging from 0.126– 1.41.

In Experiments 2 and 3, following habituation focal males were exposed to a social stimulus through a clear acrylic divider for 1 h (Fig. 1). Behavior was recorded for 1 h using a digital video surveillance system (Alibi) on Day 1 through 4 for Experiment 1 and Day 1, 4, and 5 for Experiments 2 and 3. We scored the behavior of each focal male using the BORIS event recording software (Friard and Gamba 2016). Behavior was quantified during a 10 min window 20 min after the removal of the opaque divider (i.e. 20-30 min) as this time window has been shown to adequately capture variability in behavior (O’Connell et al., 2013). In Experiment 2, we additionally scored 15-25-and 35-45 min post-divider removal to assess temporal variation in aggression. Behavior scored includes aggressive displays (lateral display, border conflict, frontal display), approaches to the shoal, and courtship displays (leads and quivers) (Supplemental Table 1). For behavior data, lateral displays, border conflicts and frontal displays were summed into aggressive displays. Courtship displays and approaches to the shoal were rare and thus not included in subsequent analyses.

### Experiment 1: Habituation Manipulation

Males were acclimated as described above and then experienced a 4-day habituation period where i) the same (familiar) neighbor was used across all 4 days and switched compartments daily or ii) a new (unfamiliar) neighbor was used across all 4 days and moved compartments daily (Fig. 1 Experiment 1).

### Experiment 2: Context Shift

For these trials, males were acclimated and habituated as described above. 16–18 h after habituation, focal males were exposed to 1 of 3 different social stimuli: i) the same male they were habituated to in the same aquarium location (‘same neighbor-same location’) ii) the same male they were habituated to in a new aquarium location (‘same neighbor-new location’) or iii) a new male in the same location of the habituated male (‘new neighbor-same location’) (Fig. 1 Experiment 2).

### Hormone Assay

Following the 1 h stimulus exposure, focal males were removed from experimental tanks, measured and weighed, and then blood was collected through the dorsal aorta using heparinized 26-G butterfly infusion sets (SURFLO Winged Infusion Sets, Terumo). Blood was spun down in a centrifuge at 3500 rpm for 15 min. Plasma was then separated and stored at −80°C. Males were killed by rapid cervical transection, and brains were dissected for a separate study. Plasma testosterone was quantified using a competitive assay (Cayman Chemicals, Ann Arbor, MI, USA) in accordance with the manufacturer’s instructions. While 11-ketotestosterone (11-KT) is the major androgen controlling male reproduction in many teleost fish (Borg 1994), circulating levels of 11-KT are much lower and more variable than those of testosterone in haplochromine cichlids including *A. burtoni* (Dijkstra et al. 2012; Kidd et al. 2010). In our study, the average intra-assay coefficient of variation (CV) was 15.36%.

### Experiment 3: Cue Competition

Males were acclimated and habituated as described above. 16–18 h after habituation, focal males were exposed to both a new male in the same location (‘new neighbor-same location’) and the same male in a new location (‘same neighbor-new location’) presented simultaneously (Fig. 1 Experiment 3).

### Data Analysis

We performed all data analysis in R (R Core Team 2025) v. 4.5.0. For comparisons of continuous data subset by categorical variables, we conducted a paired *t*-test, Welch’s *t*-test, one-way ANOVA, or Kruskal-Wallis test as appropriate based on the number of categories and normality (determined by a Shapiro-Wilk normality test), implemented in the “stats” package. To examine associations between continuous variables, we fitted linear regression models using the “stats” package. To account for repeated measures and test multivariate effects on focal male aggression, we fitted linear mixed-effects models using “lme4” and generalized linear mixed-effects models using “glmmTMB” where error distributions required non-Gaussian families. Fixed-effects in mixed-models were evaluated using Type III analyses of deviance (Wald χ^2^ tests) implemented in “car” package. Unless otherwise stated, categorical predictors were treated as factors.

<u>Experiment 1:</u> To test whether aggression differed across days (1-4) and between treatments (familiar vs. unfamiliar), we fitted a linear mixed-effects model with Day (categorical), Treatment (categorical), and their interaction as fixed effects with individual included as a random intercept to account for repeated measures. Additionally, to examine the relationship between aggression and days, we carried out a linear regression analysis for both treatment groups. <u>Experiment 2</u>: To compare focal male aggression on day 1 vs. day 4 of habituation, we conducted a paired samples *t*-test. Next, to compare focal male aggression across stimulus treatments, we conducted a Kruskal-Wallis test. To compare testosterone levels across stimulus treatments we carried out a one-way ANOVA. We also used a Pearson correlation analysis to examine the relationship between testosterone levels and aggressive displays. Additionally, to examine aggression across different time windows, we performed a negative binomial linear mixed-effect model with time and treatment as fixed effects and individual identity and round as random intercepts. Patterns of aggression and hormones across treatments and time windows were also examined using Pearson correlations. To evaluate whether focal male aggression was predicted by male coloration and morphological traits, we fitted negative binomial generalized linear mixed-effects models (family=nbinom2, loglink) with focal male color, neighbor male color, focal male GSI, and standard length (SL) difference as fixed effects, and round as random intercepts. <u>Experiment 3:</u> We conducted a paired samples *t*-test to compare focal male aggression toward the new neighbor-same location and the same neighbor-new location stimuli when presented simultaneously. Additionally, we conducted a Welch’s *t*-test to compare focal male aggression toward the same neighbor-new location when it was presented with the new neighbor-same location simultaneously vs. by itself.

## RESULTS

### Males do not habituate to familiar neighbors in changing contexts

In Experiment 1, we tested whether focal males would habituate to familiar neighbors across 4 days if neighbors moved location. We employed a linear mixed-effects model to understand which variables affected aggression while accounting for the effect of individuals. We did not find a significant effect of day (χ^2^ = 0.31, df = 3, *p* = 0.96), treatment (χ^2^ = 1.61, df = 1, *p* = 0.20), or the interaction of day and treatment (χ^2^ = 3.41, df = 3, *p* = 0.33; Fig. 2A, Supplemental Table 2). Additionally, we did not find a significant relationship between focal male aggression and days for either the familiar treatment group (R = −0.150, *p* = 0.48, Fig. 2B) or the unfamiliar treatment group (R = 0.074, *p* = 0.76, Fig. 2B).

**Figure 2:**
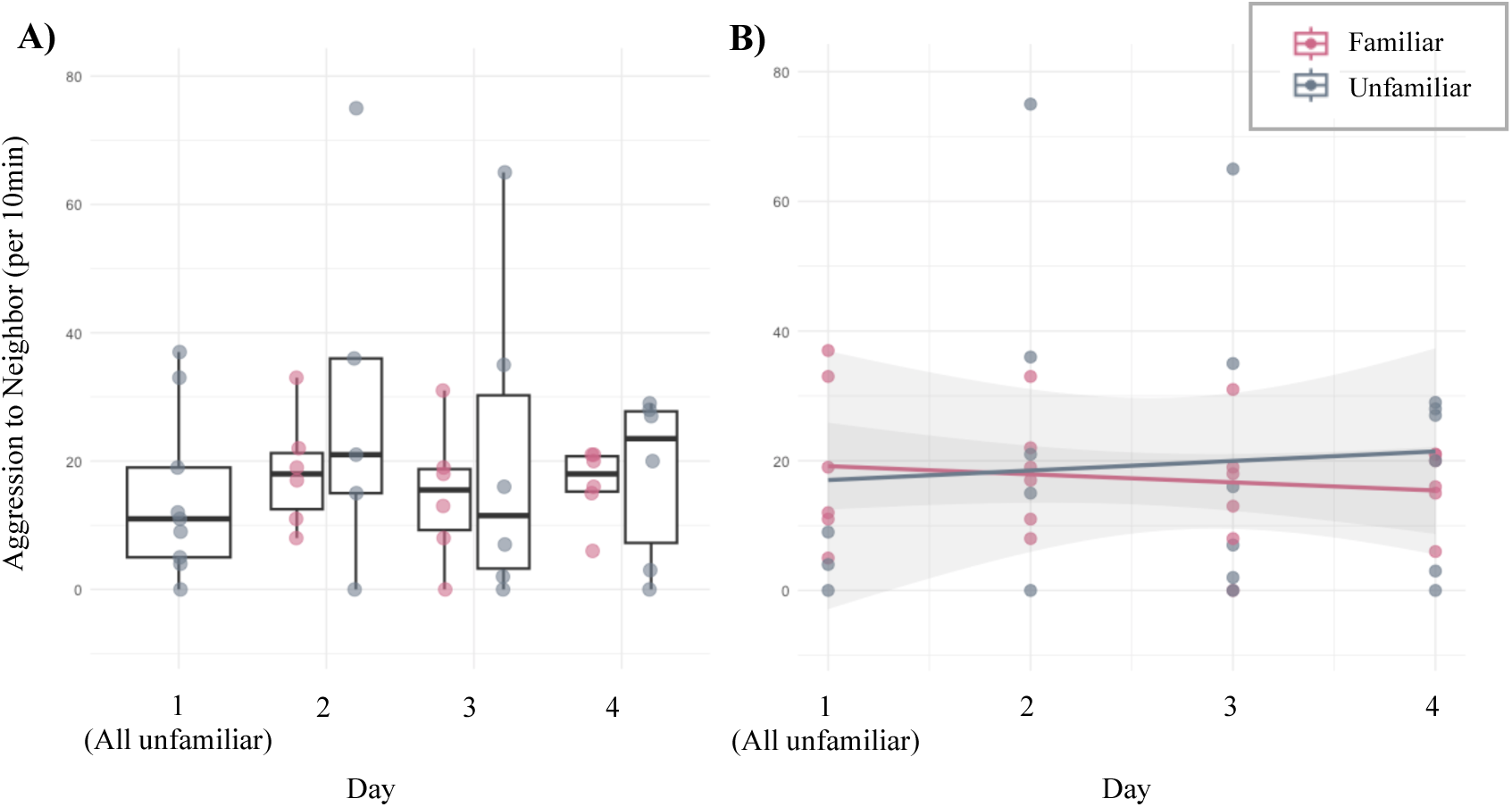
Experiment 1- Focal male aggression across 4-day habituation. (A) Focal males do not habituate to either familiar or unfamiliar neighbors across a 4-day habituation period when the neighbors are moved daily. (B) There is no relationship between aggression and day for either familiar or unfamiliar treatment.

### Dominant *A. burtoni* males show ‘Dear Enemy’ reduction in aggression to familiar neighbors

In Experiment 2, we first examined whether focal males habituate to neighboring males across 4 days. As expected, we found a significant decrease in aggression between day 1 and day 4 (t = 3.395, df = 58.17, *p* = 0.001; Fig. 3A).

**Figure 3:**
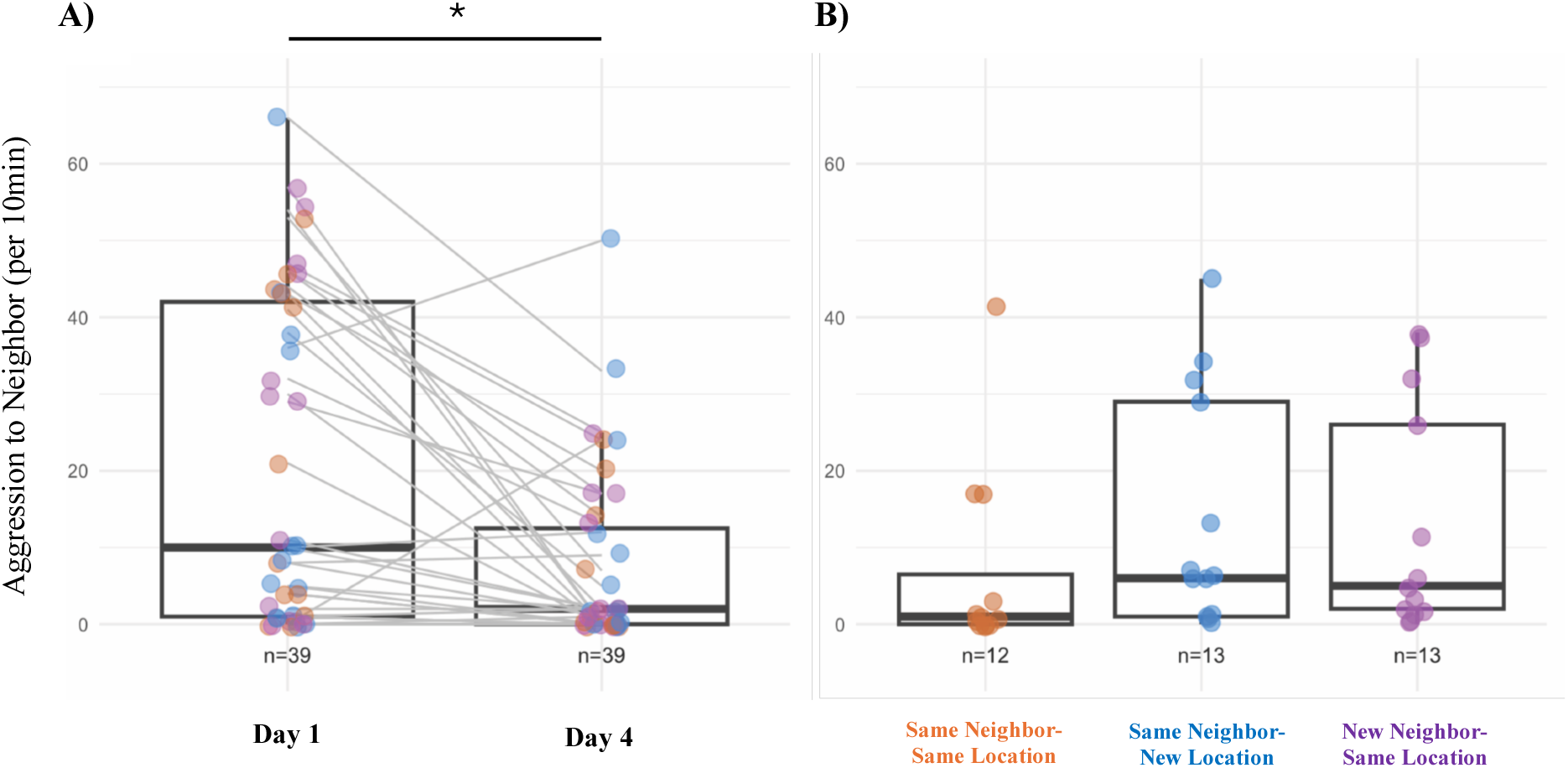
Experiment 2- Neighbor directed aggression across all treatments. (A) Aggression decreased day 1 of habituation compared to day 4. Gray lines connect the same males on day 1 and day 4. (B) Aggression across all stimulus treatments showed no significant differences. Individual data points are plotted as filled circles and colored based on the stimulus treatment experienced by each focal male. Asterisks represent significant differences (*p* < 0.05).

### Dominant males display similar levels of aggression and testosterone across treatments

We next compared levels of aggression across all stimulus treatments (same neighbor-same location, same neighbor-new location, and new neighbor-same location). We did not find a significant difference in aggression between any of the stimulus treatments (Kruskal-Wallis test: χ^2^ = 4.121, df = 2, *p* = 0.1274; Fig. 3B). Aggression levels during stimulus presentations were also similar to the reduced aggression observed on day 4 of habituation (Supplementary Fig. 3). We also compared circulating testosterone levels across stimulus treatments and did not find a significant difference between any of the groups (ANOVA: F_2,16_ = 0.640, *p* = 0.540; Fig. 4A). Note, however, that the testosterone levels we measured were higher than baseline (no stimulus) levels but lower than those observed in response to a novel intruder male, based on our previous studies (O’Connell et al., 2013; Weitekamp et al., 2017b, 2017c). We did find a significant positive correlation between testosterone levels and aggressive displays, however (R = 0.48, *p* = 0.043; Fig. 4B). To investigate whether aggressive responses changed over the course of the trial, we examined aggression across multiple time windows. Aggression showed a marginal effect of time but did not differ among treatments across time (Supplemental Fig. 1; Supplemental Table 3). Additionally, testosterone levels and aggressive behavior were also positively correlated during the first 5-15min (R = 0.49, *p* = 0.048, Supplemental Fig. 2A) and during the 35-45min time window (R = 0.69, *p* = 0.001, Supplemental Fig. 2B). We also fitted negative binomial generalized linear mixed-effects models to examine predictors of focal male aggression. Neighbor male coloration significantly predicted aggression (χ^2^ = 7.50, df=2, *p* = 0.024; Supplemental Table 4), whereas focal male coloration, focal male GSI, SL difference, and focal × neighbor color interaction did not. Parameter estimates indicated that aggression toward yellow neighbors was significantly reduced relative to blue neighbors (Supplemental Table 5).

**Figure 4:**
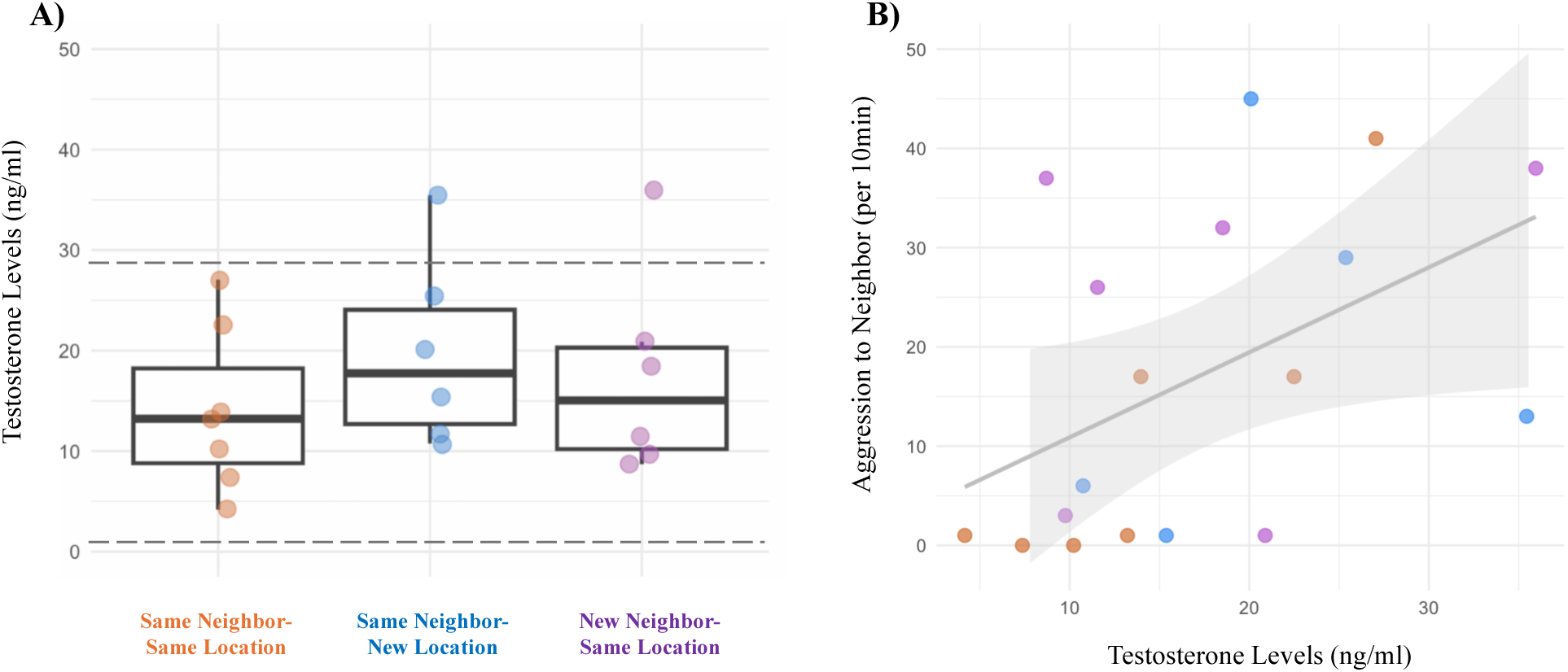
Experiment 2-Patterns of aggression and hormones across treatments. (A) Testosterone levels across stimulus treatments showed no significant differences. (B) Testosterone levels and aggressive behavior were positively correlated. Individual data points are plotted as filled circles and colored based on the stimulus treatment experienced by each focal male. Dashed lines indicate circulating levels of testosterone in territorial males with a novel intruder stimulus and no stimulus, respectively (from O’Connell et al., 2013; Weitekamp et al., 2017b, 2017c).

### Dominant males distinguish between familiar vs. unfamiliar neighbors when they are presented simultaneously

In Experiment 3, we tested whether males show different levels of aggression to the same neighbor in a new location and a new neighbor in the same location treatments when presented simultaneously. After males reduce aggression during a four-day habituation (t = 6.270, df = 16.994, *p* < 0.001, Supplementary Fig. 4), we found they aggress more to new neighbors in the same location (t = −2.224, df = 13.456, *p* = 0.0438, Fig. 5A). Additionally, we examined whether focal males showed differential levels of aggression to neighbor males when they are presented by themselves, as a single stimulus vs. when they are presented simultaneously with another stimulus neighbor. We find focal males aggress less to the same neighbor-new location male when the new neighbor-same location male is present compared to when same neighbor-new location male is presented alone (t = 2.778, df = 12.51, *p* = 0.0162; Fig. 5B). However, there is no difference in aggression when comparing the unfamiliar neighbors. Focal males showed similar levels of aggression to a new neighbor-same location neighbor presented alone compared to with another stimulus neighbor (t =-1.388, df =16.544, *p* = 0.1835; Fig. 5C).

**Figure 5:**
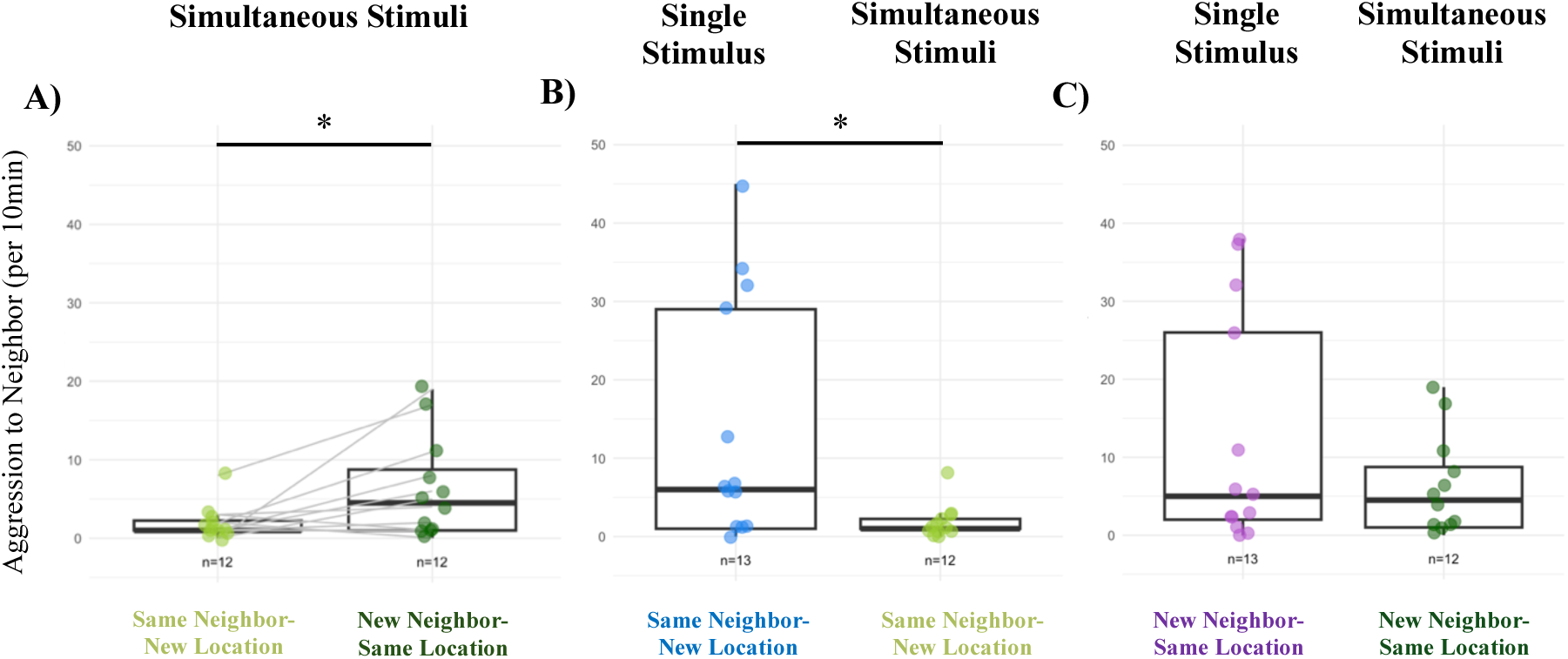
Experiment 3- Focal male aggression to neighbors when presented simultaneously. (A) Males display less aggression to a familiar neighbor in a new location when an unfamiliar neighbor is also present. (B) Further, males display less aggression to a familiar neighbor in a new location when an unfamiliar neighbor is present compared to in single stimulus treatments (i.e. Experiment 2) to the same stimulus. (C) There is no difference in focal male aggression to unfamiliar neighbors in the single stimulus treatment compared to the simultaneous stimuli treatment. Asterisks represent significant differences (*p* < 0.05).

## DISCUSSION

In the present study, we demonstrated that dominant *A. burtoni* males integrate both identity and context cues when evaluating neighbors, and that these cues jointly mediate territorial aggression. While dominant males typically habituate to neighbors across a 4-day habituation period (Weitekamp and Hofmann 2017a), we found that males no longer reduce aggression to familiar neighbors when they are located in ‘incorrect’ or novel territories, indicating spatial context is an important component of recognition. Additionally, we found that males use identity cues to distinguish among rivals, as they differentiate between a familiar neighbor in a new location and an unfamiliar neighbor. To further investigate cues involved in recognition, we examined temporal dynamics of aggression as well as male coloration patterns as possible drivers of aggressive responses. Our analyses showed slight temporal variation in aggression and suggested that neighbor coloration may influence aggressive responses. Future studies manipulating color combinations directly will help determine whether coloration functions as a reliable identity cue or signal of competitive ability. Together, these results demonstrate that habituation to rivals depends on the ecological and spatial contexts in which interactions occur, rather than a fixed behavioral rule. Our findings expand upon previous studies on animal recognition by demonstrating that individual recognition is supported by the integration of multiple information streams, including identity, spatial context, and prior social interactions, rather than reliance on a single cue (Kohda et al., 2015; Tibbetts & Dale, 2007; Wiley, 2013). These results further our understanding of the mechanisms animals use to mediate territorial aggression and maintain stable communities.

### *A. burtoni* males rely on context cues for social habituation

We found that focal males did not habituate to familiar neighbors if they moved location during the habituation period. In fact, males aggressed towards these familiar males at levels similar as towards unfamiliar males. However, when focal males are given visual exposure to the same neighbor in the same location across a 4-day habituation period, we found they do reduce aggression and establish a ‘dear enemy’ relationship, which is consistent with previous work in *A. burtoni* (Weitekamp and Hofmann 2017a) as well as other cichlid species (e.g. *Oreochromis mossambicus*: Aires et al. 2015; *Amatitlania nigrofasciata*: Leiser and Itzkowitz 1999; *N. pulcher*: Frostman and Sherman 2004). The fact that males did not habituate to familiar neighbors when spatial cues were disrupted suggests a reliance on a place-identity associative memory. Similar forms of context-dependent recognition are found across animals where spatial or environmental cues modulate behavior even after neighbor identity has been learned. For example, territorial songbirds rely on territorial boundary locations as part of neighbor identity (Godard 1991; Akcay et al. 2010), and social insects integrate spatial and chemical information to regulate nestmate recognition (Buczkowski & Silverman, 2005; Sturgis & Gordon, 2011). This suggests that individual recognition is not solely a property of the receiver’s memory, but of how memories are embedded within stable environmental contexts.

Future research testing on how spatial associations interact with identity cues across species with different territorial systems will help clarify whether place-identity coupling is a general feature of recognition systems or one shaped by specific ecological demands. Additionally, experimentally manipulating identity and context cues independently would help disentangle the relative contributions of spatial stability, cue modality, and learning to the recognition processes revealed here.

### Focal males display differential levels of aggression to territorial challenges based on the identity of neighbors

To investigate whether *A. burtoni* males use individually distinctive features for recognition, we measured aggression toward neighbors that varied in identity and location. In single stimulus treatments, males showed similar aggression and testosterone levels across treatments, closely matching the reduced aggression observed in Day 4 of habituation. This pattern suggests that repeated exposure may reduce threat perception, consistent with general principles of habituation (Blumstein, 2016; Raderschall et al., 2011). Because *A. burtoni* males exhibited the dear enemy effect, a form of social habituation, in response to repeated exposure to intruder males, they could be maintaining low levels of aggression during experimental treatments despite neighbor location and/or identity changing. Although testosterone did not differ among treatments, circulating levels remained elevated relative to levels reported for unstimulated territorial males while remaining lower than those reported for territorial males exposed to a novel intruder (O’Connell et al., 2013; Weitekamp et al., 2017b, 2017c). Together, these findings suggest that focal males continued to perceive neighboring males as socially salient, but no longer as high-threat competitors as indicated by their reduced aggressive and endocrine responses.

Features of a stimulus (e.g. intensity, frequency, modality, duration, novelty) can also influence its perceived threat or saliency and impact the timing of habituation and sensitization processes and/or if they occur. For example, animals habituate slower to high-intensity cues such as predators, attractive mates, or rivals (Canty & Gould, 1995; Jenkins & Rowland, 2000; Rowland, 2000). Thus, individuals must balance conserving energy through habituation with the risk of under-responding to genuinely threatening intruders (Jenkins & Rowland, 2000; Peeke & Figler, 1997). The rate or time course of habituation could therefore be very important especially in complex environments where animals are exposed to multiple stimuli and it is advantageous to modulate behavior to different stimuli over relatively short timescales. Work in *N. pulcher* demonstrates time-dependent declines in aggression and suggests that familiar individuals appearing in unexpected locations trigger transient increases in aggression (Saeki et al., 2018). To investigate whether aggressive responses changed over the course of the trial, we examined aggression across multiple time windows. Aggression showed temporal variation but did not differ among treatments across time. These patterns suggest a general temporal shift in aggression, with only short-lived differences among groups.

More broadly, these patterns align with recognition and threat-assessment mechanisms found across organisms where they must integrate multiple cues (e.g. identity, appearance, spatial location, and interaction history) to regulate aggression and minimize costs of agonistic interactions. For example, birds depend on both identity cues and territory boundaries for neighbor-stranger discrimination (Godard 1991; Akcay et al. 2010); lizards use color signals, spatial familiarity, and prior interaction history to modulate aggression (Carazo et al., 2008; Whiting et al., 2006); social insects integrate chemical identity cues with spatial context during nestmate recognition (Buczkowski & Silverman, 2005; Sturgis & Gordon, 2011); and mammals rely on multimodal cues and previous experience to assess rivals and regulate territorial behavior (e.g. primates and ground squirrels: Hare and Atkins 2001; Cheney and Seyfarth 2008). Future work that examines how animals weigh and integrate multiple social cues during rival assessment will help reveal the decision rules underlying recognition across taxa.

### Males differentially aggress towards familiar and unfamiliar neighbors when presented simultaneously

To test whether males can distinguish and/or differentially respond to familiar males in a new location and unfamiliar males, we presented focal males with both stimuli simultaneously. We found that males differentially respond to these neighbor types and aggress less to a familiar neighbor in a new location compared to an unfamiliar neighbor. This finding is consistent with the threat-level hypothesis and suggests that focal males are adjusting their aggression in response to the perceived threat level of their neighbors. Similarly, when comparing aggression towards the familiar neighbor in a new location in the single vs. simultaneous stimulus treatments, males showed lower aggression in the simultaneous treatment, suggesting that familiar males are evaluated as less threatening in comparison to a higher-risk rival. More generally, such competitive evaluation mirrors decision rules found across animals where individuals integrate identity, context and perceived threat when choosing whom to aggress, avoid, or court (Akcay et al., 2010; Carazo et al., 2008; Cheney & Seyfarth, 2008; Hyman, 2002; Sturgis & Gordon, 2011; Whiting et al., 2006). Future studies examining whether males integrate multiple cues additively or hierarchically by using conflicting information would help to reveal the rules of cue weighing animals may be using.

## CONCLUSIONS

*A. burtoni* males live in dense communities in their natural habitats where repeated interactions with the same individuals make recognition mechanisms advantageous to establish and maintain territories and reduce the costs of agonistic interactions. Our results show that *A. burtoni* males rely on both identity and context cues to regulate territorial aggression, and that these cues interact dynamically when individuals evaluate the relative threat posed by familiar and unfamiliar rivals. For example, territorial location, neighbor coloration, previous interactions, temporal dynamics could all be cues males are using to recognize individuals and inform decision-making. These findings highlight recognition as a multi-component process that reflects broader principles of cue integration seen across territorial animals. Future research that compares how different species encode, weight, and update identity and contextual information will help establish the generality of these decision rules and the evolutionary pressures that shape recognition systems in social and territorial species.

## Ethics

This work was approved by the Institutional Animal Care and Use Committee at the University of Texas at Austin. Clinical trial number: not applicable.

## References

Aires, R. F., Oliveira, G. A., Oliveira, T. F., Ros, A. F. H., & Oliveira, R. F. (2015). Dear Enemies Elicit Lower Androgen Responses to Territorial Challenges than Unfamiliar Intruders in a Cichlid Fish. PLOS ONE, 10(9), e0137705. 10.1371/journal.pone.0137705

Akcay, C., Reed, V. A., Campbell, S. E., Templeton, C. N., & Beecher, M. D. (2010). Indirect reciprocity: Song sparrows distrust aggressive neighbours based on eavesdropping. Animal Behaviour, 80(6), 1041–1047. 10.1016/j.anbehav.2010.09.009

Akçay, Ç., Wood, W. E., Searcy, W. A., Templeton, C. N., Campbell, S. E., & Beecher, M. D. (2009). Good neighbour, bad neighbour: Song sparrows retaliate against aggressive rivals. Animal Behaviour, 78(1), 97–102. 10.1016/j.anbehav.2009.03.023

Arnold, B. D., & Wilkinson, G. S. (2011). Individual specific contact calls of pallid bats (Antrozous pallidus) attract conspecifics at roosting sites. Behavioral Ecology and Sociobiology, 65(8), 1581–1593. 10.1007/s00265-011-1168-4

Aubin, T., Jouventin, P., & Hildebrand, C. (2000). Penguins use the two–voice system to recognize each other. Proceedings of the Royal Society of London. Series B: Biological Sciences. (world). 10.1098/rspb.2000.1112

Backwell, P. R. Y., & Jennions, M. D. (2004). Coalition among male fiddler crabs. Nature, 430(6998), 417–417. 10.1038/430417a

Bee, M. A., & Gerhardt, H. C. (2002). Individual voice recognition in a territorial frog (Rana catesbeiana). Proceedings of the Royal Society B: Biological Sciences, 269(1499), 1443–1448. 10.1098/rspb.2002.2041

Beecher, M. D. (1989). Signalling systems for individual recognition: An information theory approach. Animal Behaviour, 38(2), 248–261. 10.1016/S0003-3472(89)80087-9

Blumstein, D. T. (2016). Habituation and sensitization: New thoughts about old ideas. Animal Behaviour, 120, 255–262. 10.1016/j.anbehav.2016.05.012

Bonnie, K. E., & Earley, R. L. (2007). Expanding the scope for social information use. Animal Behaviour, 74(2), 171–181. 10.1016/j.anbehav.2006.12.009

Briseño-Jaramillo, M., Estrada, A., & Lemasson, A. (2015). Individual voice recognition and an auditory map of neighbours in free-ranging black howler monkeys (Alouatta pigra). Behavioral Ecology and Sociobiology, 69(1), 13–25. 10.1007/s00265-014-1813-9

Buczkowski, G., & Silverman, J. (2005). Context-dependent nestmate discrimination and the effect of action thresholds on exogenous cue recognition in the Argentine ant. Animal Behaviour, 69(3), 741–749. 10.1016/j.anbehav.2004.06.027

Canty, N., & Gould, J. L. (1995). The hawk/goose experiment: Sources of variability. Animal Behaviour, 50(4), 1091–1095. 10.1016/0003-3472(95)80108-1

Carazo, P., Font, E., & Desfilis, E. (2008). Beyond ‘nasty neighbours’ and ‘dear enemies’? Individual recognition by scent marks in a lizard (Podarcis hispanica). Animal Behaviour, 76(6), 1953–1963. 10.1016/j.anbehav.2008.08.018

Carlson, N. V., Kelly, E. M., & Couzin, I. (2020). Individual vocal recognition across taxa: A review of the literature and a look into the future. Philosophical Transactions of the Royal Society B: Biological Sciences, 375(1802), 20190479. 10.1098/rstb.2019.0479

Cheney, D. L., & Seyfarth, R. M. (2008). Baboon Metaphysics: The Evolution of a Social Mind. University of Chicago Press. https://press.uchicago.edu/ucp/books/book/chicago/B/bo5387727.html

Chuang, M.-F., Kam, Y.-C., & Bee, M. A. (2017). Territorial olive frogs display lower aggression towards neighbours than strangers based on individual vocal signatures. Animal Behaviour, 123, 217–228. 10.1016/j.anbehav.2016.11.001

Dale, J. (2006). Intraspecific variation in coloration (Vol. 2). University Press. https://books.google.com/books/about/Bird_Coloration.html?id=BnM8musSTS4C

Dale, J., B. Lank, D., & Kern Reeve, H. (2001). Signaling Individual Identity versus Quality: A Model and Case Studies with Ruffs, Queleas, and House Finches. The American Naturalist. (world). 10.1086/320861

Desjardins, J., Hofmann, H., & Fernald, F. (2012). Social context influences aggressive and courtship behavior in a cichlid fish. PloS One, 7(7). 10.1371/journal.pone.0032781

Dijkstra, P. D., Verzijden, M. N., Groothuis, T. G. G., & Hofmann, H. A. (2012). Divergent hormonal responses to social competition in closely related species of haplochromine cichlid fish. Hormones and Behavior, 61(4), 518–526. 10.1016/j.yhbeh.2012.01.011

Douglas, R., & Djamgoz, M. (2012). The Visual System of Fish. Springer Science & Business Media.

Espmark, Y. (1971). Individual Recognition By Voice in Reindeer Mother-Young Relationship. Field Observations and Playback Experiments. 10.1163/156853971X00438

Fernald, R. D. (1977). Quantitative behavioural observations of *Haplochromis burtoni* under semi-natural conditions. Animal Behaviour, 25, 643–653. 10.1016/0003-3472(77)90115-4

Fernald, R. D., & Hirata, N. R. (1977). Field study of *Haplochromis burtoni*: Quantitative behavioural observations. Animal Behaviour, 25, 964–975. 10.1016/0003-3472(77)90048-3

Fleischer, J., & Krieger, J. (2018). Insect Pheromone Receptors – Key Elements in Sensing Intraspecific Chemical Signals. Frontiers in Cellular Neuroscience, 12. 10.3389/fncel.2018.00425

Friard, O., & Gamba, M. (2016). BORIS: A free, versatile open-source event-logging software for video/audio coding and live observations. Methods in Ecology and Evolution, 7(11), 1325–1330. 10.1111/2041-210X.12584

Frostman, P., & Sherman, P. T. (2004). Behavioral response to familiar and unfamiliar neighbors in a territorial cichlid, Neolamprologus pulcher. Ichthyological Research, 51(3), 283–285. 10.1007/s10228-004-0223-9

Gentner, T. Q., & Hulse, S. H. (1998). Perceptual mechanisms for individual vocal recognition in European starlings,*Sturnus vulgaris*. Animal Behaviour, 56(3), 579–594. 10.1006/anbe.1998.0810

Gleason, E. D., Fuxjager, M. J., Oyegbile, T. O., & Marler, C. A. (2009). Testosterone release and social context: When it occurs and why. *Frontiers in Neuroendocrinology*, Hormones & Social Behavior, 30(4), 460–469. 10.1016/j.yfrne.2009.04.009

Godard, R. (1991). Long-term memory of individual neighbours in a migratory songbird. Nature, 350(6315), 228–229. 10.1038/350228a0

Goymann, W., Moore, I. T., & Oliveira, R. F. (2019). Challenge Hypothesis 2.0: A Fresh Look at an Established Idea. BioScience, 69(6), 432–442. 10.1093/biosci/biz041

Grosenick, L., Clement, T. S., & Fernald, R. D. (2007). Fish can infer social rank by observation alone. Nature, 445(7126), 429–432. 10.1038/nature05511

Guilford, T., & Dawkins, M. S. (1995). What are conventional signals? Animal Behaviour, 49(6), 1689–1695. 10.1016/0003-3472(95)90090-X

Hare, J. F., & Atkins, B. A. (2001). The squirrel that cried wolf: Reliability detection by juvenile Richardson’s ground squirrels (Spermophilus richardsonii). Behavioral Ecology and Sociobiology, 51(1), 108–112. 10.1007/s002650100414

Hebets, E. A., & Papaj, D. R. (2005). Complex signal function: Developing a framework of testable hypotheses. Behavioral Ecology and Sociobiology, 57(3), 197–214. 10.1007/s00265-004-0865-7

Hofmann, H. A., & Schildberger, K. (2001). Assessment of strength and willingness to fight during aggressive encounters in crickets. Animal Behaviour, 62(2), 337–348. 10.1006/anbe.2001.1746

Hyman, J. (2002). Conditional strategies in territorial defense: Do Carolina wrens play tit-for-tat? Behavioral Ecology, 13(5), 664–669. 10.1093/beheco/13.5.664

Jenkins, J. R., & Rowland, W. J. (2000). Stimulus-specific and response-specific habituation in courting stickleback: Developmental and functional considerations. Behaviour, 137(7–8), 933–945. 10.1163/156853900502547

Johnstone, R. A. (1997). Recognition and the evolution of distinctive signatures: When does it pay to reveal identity? Proceedings of the Royal Society B: Biological Sciences, 264(1387), 1547–1553. 10.1098/rspb.1997.0215

Kendrick, K. M., Atkins, K., Hinton, M. R., Heavens, P., & Keverne, B. (1996). Are faces special for sheep? Evidence from facial and object discrimination learning tests showing effects of inversion and social familiarity. Behavioural Processes, 38(1), 19–35. 10.1016/0376-6357(96)00006-X

Kidd, C. E., Kidd, M. R., & Hofmann, H. A. (2010). Measuring multiple hormones from a single water sample using enzyme immunoassays. General and Comparative Endocrinology, 165(2), 277–285. 10.1016/j.ygcen.2009.07.008

Kohda, M., Jordan, L. A., Hotta, T., Kosaka, N., Karino, K., Tanaka, H., Taniyama, M., & Takeyama, T. (2015). Facial Recognition in a Group-Living Cichlid Fish. PLOS ONE, 10(11), e0142552. 10.1371/journal.pone.0142552

Leiser, J., & Itzkowitz, M. (1999). The Benefits of Dear Enemy Recognition in Three-Contender Convict Cichlid (Cichlasoma Nigrofasciatum) Contests. 10.1163/156853999501685

Lesbarrères, D., & Lodé, T. (2002). Variations in male calls and responses to an unfamiliar advertisement call in a territorial breeding anuran, Rana dalmatina: Evidence for a “dear enemy” effect. Ethology Ecology & Evolution, 14(4), 287–295. 10.1080/08927014.2002.9522731

Marler, C. A., Oyegbile, T. O., Plavicki, J., & Trainor, B. C. (2005a). Response to Wingfield’s commentary on “A continuing saga: The role of testosterone in aggression.” Hormones and Behavior, 48(3), 256–258. 10.1016/j.yhbeh.2005.05.010

Martin-Malivel, J., & Okada, K. (2007). Human and chimpanzee face recognition in chimpanzees (Pan troglodytes): Role of exposure and impact on categorical perception. Behavioral Neuroscience, 121(6), 1145–1155. 10.1037/0735-7044.121.6.1145

Maruska, K. P. (2014). Social regulation of reproduction in male cichlid fishes. General and Comparative Endocrinology, Perspectives in Cichlid Endocrinology, 207, 2–12. 10.1016/j.ygcen.2014.04.038

Maruska, K. P., Becker, L., Neboori, A., & Fernald, R. D. (2013). Social descent with territory loss causes rapid behavioral, endocrine and transcriptional changes in the brain. Journal of Experimental Biology, 216(19), 3656–3666. 10.1242/jeb.088617

Maruska, K. P., & Fernald, R. D. (2018). Astatotilapia burtoni: A Model System for Analyzing the Neurobiology of Behavior. ACS Chemical Neuroscience, 9(8), 1951–1962. 10.1021/acschemneuro.7b00496

McLennan, D. A. (2004). Male Brook Sticklebacks’ (Culaea inconstans) Response to Olfactory Cues. Behaviour, 141(11/12), 1411–1424. http://www.jstor.org/stable/4536213

Müller, C. A., & Manser, M. B. (2007). ‘Nasty neighbours’ rather than ‘dear enemies’ in a social carnivore. Proceedings of the Royal Society B: Biological Sciences, 274(1612), 959. 10.1098/rspb.2006.0222

Myrberg, A. A., & Riggio, R. J. (1985). Acoustically mediated individual recognition by a coral reef fish (*Pomacentrus partitus*). Animal Behaviour, 33(2), 411–416. 10.1016/S0003-3472(85)80065-8

O’Connell, L. A., Rigney, M. M., Dykstra, D. W., & Hofmann, H. A. (2013). Neuroendocrine Mechanisms Underlying Sensory Integration of Social Signals. Journal of Neuroendocrinology, 25(7), 644–654. 10.1111/jne.12045

Olendorf, R., Getty, T., Scribner, K., & Robinson, S. K. (2004). Male red–winged blackbirds distrust unreliable and sexually attractive neighbours. Proceedings of the Royal Society of London. Series B: Biological Sciences, 271(1543), 1033–1038. 10.1098/rspb.2004.2687

Peake, T. M. (2005). Eavesdropping in communication networks. In P. K. McGregor (Ed.), Animal Communication Networks (pp. 13–37). Cambridge University Press. 10.1017/CBO9780511610363.004

Peeke, H. V. S., & Figler, M. H. (1997). Form and Function of Habituation and Sensitization of Male Courtship in the Three-Spined Stickleback (Gasterosteus aculeatus L.). Behaviour, 134(15/16), 1273–1287. https://www.jstor.org/stable/4535504

Raderschall, C. A., Magrath, R. D., & Hemmi, J. M. (2011). Habituation under natural conditions: Model predators are distinguished by approach direction. Journal of Experimental Biology, 214(24), 4209–4216. 10.1242/jeb.061614

Robertson, B. C. (1996). Vocal mate recognition in a monogamous, flock-forming bird, the silvereye, Zosterops lateralis. Animal Behaviour, 51(2), 303–311. 10.1006/anbe.1996.0030

Rosenfeld, S. A., & Van Hoesen, G. W. (1979). Face recognition in the rhesus monkey. Neuropsychologia, 17(5), 503–509. 10.1016/0028-3932(79)90057-5

Rowland, W. J. (2000). Habituation and development of response specificity to a sign stimulus: Male preference for female courtship posture in stickleback. Animal Behaviour, 60(1), 63–68. 10.1006/anbe.2000.1462

Saeki, T., Sogawa, S., Hotta, T., & Kohda, M. (2018). Territorial fish distinguish familiar neighbours individually. Behaviour, 155(4), 279–293. (129181386). 10.1163/1568539X-00003489

Siebeck, U. E., Parker, A. N., Sprenger, D., Mäthger, L. M., & Wallis, G. (2010). A Species of Reef Fish that Uses Ultraviolet Patterns for Covert Face Recognition. Current Biology, 20(5), 407–410. 10.1016/j.cub.2009.12.047

Smadja, C., & Butlin, R. K. (2009). On the scent of speciation: The chemosensory system and its role in premating isolation. Heredity, 102(1), 77–97. 10.1038/hdy.2008.55

Sturgis, S. J., & Gordon, D. M. (2011). Nestmate recognition in ants (Hymenoptera: Formicidae): A review.

Tanaka, J., & Farah, M. (1993). Parts and wholes in face recognition. The Quarterly Journal of Experimental Psychology. A, Human Experimental Psychology, 46(2). 10.1080/14640749308401045

Temeles, E. J. (1994). The role of neighbours in territorial systems: When are they “dear enemies”? Animal Behaviour, 47(2), 339–350. 10.1006/anbe.1994.1047

Tibbetts, E. A. (2002). Visual signals of individual identity in the wasp Polistes fuscatus. Proceedings of the Royal Society of London. Series B: Biological Sciences. (world). 10.1098/rspb.2002.2031

Tibbetts, E. A. (2025). Individual face recognition in wasps. Trends in Cognitive Sciences. 10.1016/j.tics.2025.08.004

Tibbetts, E. A., & Dale, J. (2007). Individual recognition: It is good to be different. Trends in Ecology & Evolution, 22(10), 529–537. 10.1016/j.tree.2007.09.001

Tibbetts, E. A., Pardo-Sanchez, J., & Weise, C. (2022). The establishment and maintenance of dominance hierarchies. Philosophical Transactions of the Royal Society B: Biological Sciences, 377(1845), 20200450. 10.1098/rstb.2020.0450

Valone, T. J. (2007). From eavesdropping on performance to copying the behavior of others: A review of public information use. Behavioral Ecology and Sociobiology, 62(1), 1–14. 10.1007/s00265-007-0439-6

Vasconcelos, M. (2008). Transitive inference in non-human animals: An empirical and theoretical analysis. Behavioural Processes, 78(3), 313–334. 10.1016/j.beproc.2008.02.017

Weitekamp, C. A., & Hofmann, H. A. (2017a). Neuromolecular correlates of cooperation and conflict during territory defense in a cichlid fish. Hormones and Behavior, 89, 145–156. 10.1016/j.yhbeh.2017.01.001

Weitekamp, C. A., Nguyen, J., & Hofmann, H. A. (2017b). Neuromolecular Regulation of Aggression Differs by Social Role during Joint Territory Defense. Integrative and Comparative Biology, 57(3), 631–639. 10.1093/icb/icx009

Weitekamp, C. A., Nguyen, J., & Hofmann, H. A. (2017c). Social context affects behavior, preoptic area gene expression, and response to D2 receptor manipulation during territorial defense in a cichlid fish. Genes, Brain and Behavior, 16(6), 601–611. 10.1111/gbb.12389

Werba, J. A., Stuckert, A. M., Edwards, M., & McCoy, M. W. (2022). Stranger danger: A meta-analysis of the dear enemy hypothesis. Behavioural Processes, 194, 104542. 10.1016/j.beproc.2021.104542

Whiting, M. J., Stuart-Fox, D. M., O’Connor, D., Firth, D., Bennett, N. C., & Blomberg, S. P. (2006). Ultraviolet signals ultra-aggression in a lizard. Animal Behaviour, 72(2), 353–363. 10.1016/j.anbehav.2005.10.018

Wiley, R. H. (2013). Signal Detection, Noise, and the Evolution of Communication. In H. Brumm (Ed.), Animal Communication and Noise (Vol. 2, pp. 7–30). Springer Berlin Heidelberg. 10.1007/978-3-642-41494-7_2

Wingfield, J. C., Hegner, R. E., Dufty, Alfred M., & Ball, G. F. (1990). The “Challenge Hypothesis”: Theoretical Implications for Patterns of Testosterone Secretion, Mating Systems, and Breeding Strategies. The American Naturalist, 136(6), 829–846. 10.1086/285134

Wunderle, J. M., Jr. (1978). Differential Response of Territorial Yellowthroats to the Songs of Neighbors and Non-neighbors. The Auk, 95(2), 389–395. 10.1093/auk/95.2.389

Wyatt, T. (2003). Pheromones and Animal Behaviour: Communication by Smell and Taste. Cambridge University Press. https://books.google.com/books/about/Pheromones_and_Animal_Behaviour.html?id=aty3hz6rdlgC

Ydenberg, R. C., Giraldeau, L. A., & Falls, J. B. (1988). Neighbours, strangers, and the asymmetric war of attrition. Animal Behaviour, 36(2), 343–347. 10.1016/S0003-3472(88)80004-6

