## Supplementary materials for "Context and Identity Cues Mediate Individual Recognition in a Highly Social Cichlid Fish"

**Supplemental Table 1. Behavioral displays quantified for focal males.**

| **Type** | **Display** | **Description** |
| --- | --- | --- |
| **Aggressive** | Frontal display | One male aggressively displays towards another |
|  | Lateral display | Aggressive display where male presents his fins laterally with occasional quivering |
|  | Border conflict | Confrontation between two males to define territorial boundaries |
| **Reproductive** | Approach/Lead | Males swim towards females |
|  | Quiver | Male approaches female and presents fins laterally, occasionally quivering |
| **Social** | Approach | Swim quickly toward female or juvenile |

**Supplemental Table 2. Experiment 1-Linear mixed model testing the effects of treatment on aggression during habituation.** We performed a linear mixed-effects model with day (1-4) and treatment (familiar vs. unfamiliar) as fixed effects and individual as a random intercept to account for repeated measures. We did not find significant effects of day, treatment, or interaction effects.

| **Response**  **Variable** | **Predictor** | **χ^2^** | **df** | ***p*** |
| --- | --- | --- | --- | --- |
| ***Focal male aggression*** | ***Day*** *(1-4)* | 0.31 | 3 | 0.959 |
|  | ***Treatment*** (Familiar vs. Unfamiliar) | 1.61 | 1 | 0.204 |
|  | ***Day*** * ***Treatment*** | 3.41 | 3 | 0.333 |

| **Response**  **Variable** | **Predictor** | **χ^2^** | **df** | ***p*** |
| --- | --- | --- | --- | --- |
| ***Focal male aggression*** | ***Time*** | 5.98 | 2 | 0.050 |
|  | ***Treatment*** | 5.29 | 2 | 0.071 |
|  | ***Time*** * ***Treatment*** | 3.45 | 4 | 0.485 |

New Neighbor-Same Location

Same Neighbor-Same Location

Same Neighbor-New Location

**Supplemental Figure 1: Experiment 2-Male aggression to neighbor across stimulus treatments and different time windows.** Male aggression was not different among stimulus treatments or across time windows. Individual data points are plotted as filled circles and colored based on the stimulus treatment experienced by each focal male.

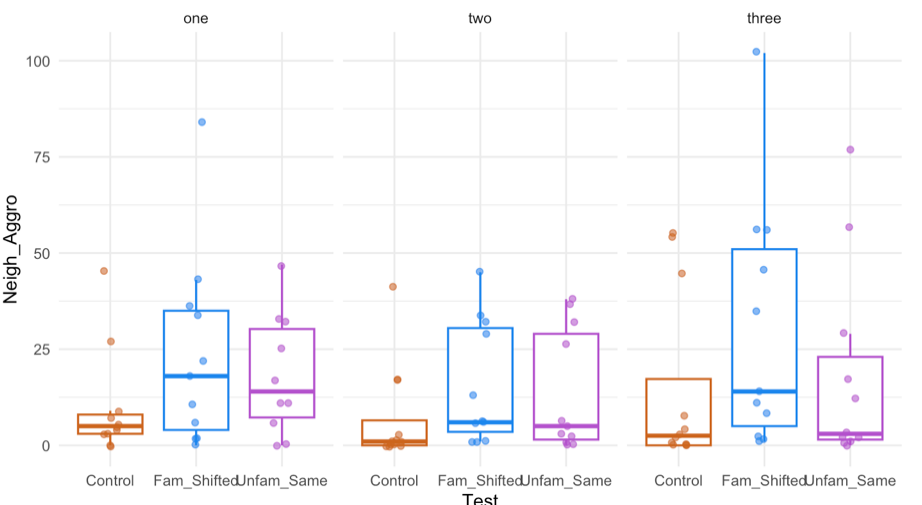

5-15

20-30

35-45

Time (minutes)

Aggression to Neighbor (per 10min)

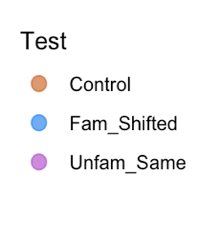

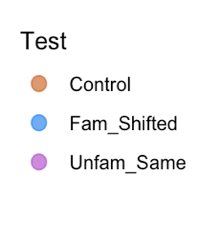

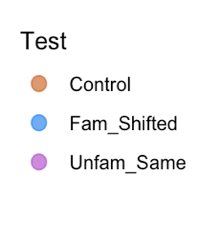

**Supplemental Table 3: Experiment 2- Effects of time window and treatment on focal male aggression.** We performed a negative binomial generalized linear mixed-effects model with time window and treatment as fixed effects and individual identity and round as random intercepts to account for repeated measures. Aggression varied across time windows, but we did not detect a significant interaction between time window and treatment.

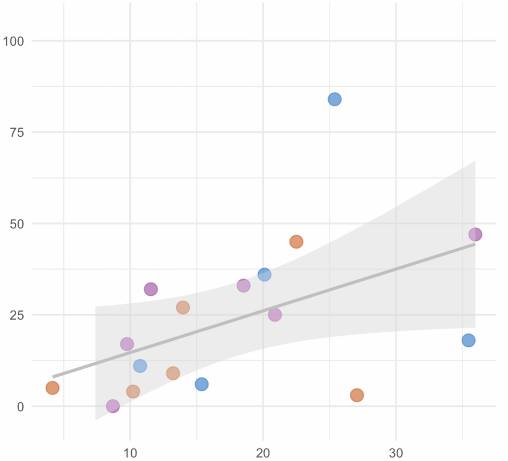

Testosterone Levels (ng/ml)

Aggression to Neighbor (per 10min)

**A)**

**Supplemental Figure 2: Experiment 2- Patterns of aggression and hormones across treatments and time windows.** Testosterone levels and aggressive behavior were positively correlated during the 5-15min time window (A; R = 0.49, *p* = 0.048) as well as the 35-45min time window (B; R = 0.69, *p* = 0.001). Individual data points are plotted as filled circles and colored based on the stimulus treatment experienced by each focal male.

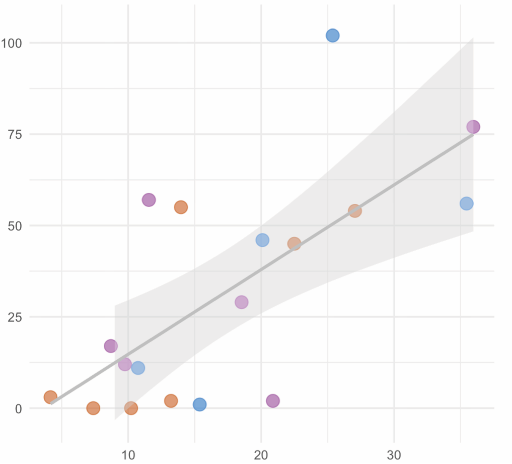

Testosterone Levels (ng/ml)

Aggression to Neighbor (per 10min)

**B)**

Same Neighbor-Same Location

Same Neighbor-New Location

New Neighbor-Same Location

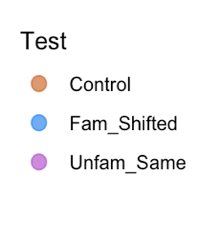

**Supplementary Table 4. Experiment 2-General linear mixed effects models testing the predictors of neighbor-directed aggression.** We performed negative binomial GLMMs to examine whether focal male color, neighbor male color, focal male GSI, and size (SL) difference between males best predicted focal male aggression. The best-fit model included additive effects of focal and neighbor color, focal male GSI, and SL difference as fixed effects with round as random effects. The inclusion of focal $\times$ neighbor color interaction did not improve model fit (LRT: χ^2^ = 5.08, df = 4, *p* = 0.28, $\Delta$AIC = -2.92). Fixed effects were evaluated using Type III Analyses of Deviance (Wald χ^2^ tests).

| **Response**  **Variable** |  | **Predictor** | χ**^2^** | **df** | ***p*-value** |
| --- | --- | --- | --- | --- | --- |
| ***Focal male aggression*** | *Model AIC = 235.79* | ***Focal male color*** | 3.8228 | 2 | 0.1479 |
|  |  | ***Neighbor color*** | 7.4962 | 2 | 0.0236 |
|  |  | ***Focal male GSI*** | 1.8186 | 1 | 0.1775 |
|  |  | ***SL diff*** | 0.9369 | 1 | 0.3331 |

**Supplementary Table 5. Parameter estimates from the best-fit negative binomial GLMM predicting neighbor-directed aggression.** Parameter estimates from the final additive negative binomial generalized linear mixed model predicting focal male aggression towards neighbors. Blue morph males were used as the reference category for focal and neighbor color effects. Positive estimates indicate increased aggression relative to the reference category, whereas negative estimates indicate reduced aggression. Experimental round was used as a random effect.

| **Predictor** | **Estimate (**$\boldsymbol{\beta)}$ | | **SE** | ***z*** | ***p-value*** | |
| --- | --- | --- | --- | --- | --- | --- |
| ***Intercept*** | 3.0733 | 0.8700 | | 3.5330 | | 0.0004 |
| ***Focal color: Blue Yellow*** | 0.8037 | 0.9626 | | 0.8350 | | 0.4037 |
| ***Focal color: Yellow*** | 1.6412 | 0.9673 | | 1.6970 | | 0.0898 |
| ***Neighbor color:***  ***Blue-Yellow*** | -0.3468 | 0.8481 | | -0.4090 | | 0.6826 |
| ***Neighbor color:***  ***Yellow*** | -1.7165 | 0.8536 | | -2.0110 | | 0.0443 |
| ***Focal male GSI*** | -1.1105 | 0.8235 | | -1.3490 | | 0.1775 |
| ***SL diff*** | -1.1709 | 1.2180 | | -0.9680 | | 0.3331 |

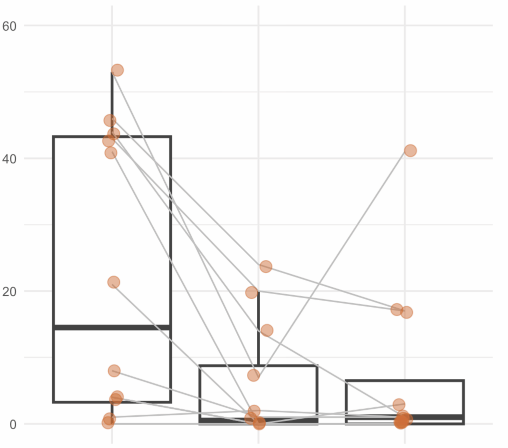

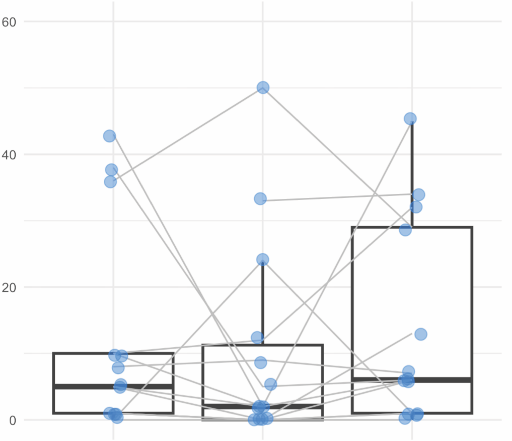

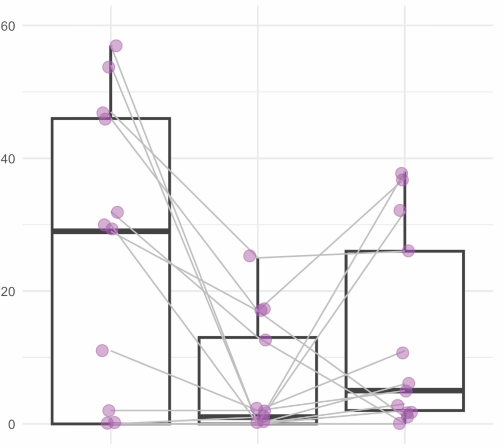

Aggression to Neighbor (per 10min)

**Habituation**

**Stimulus**

**Treatment**

**Habituation**

**Stimulus**

**Treatment**

**Habituation**

**Stimulus**

**Treatment**

**A)**

**B)**

**C)**

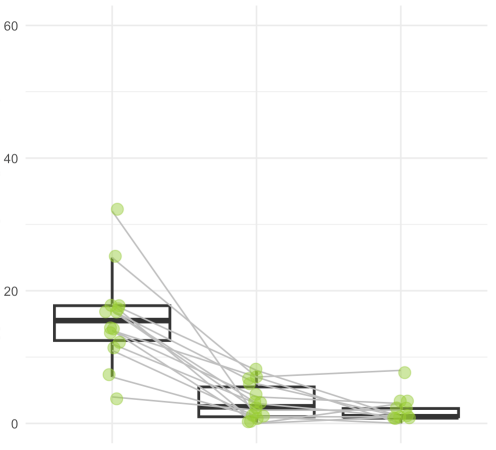

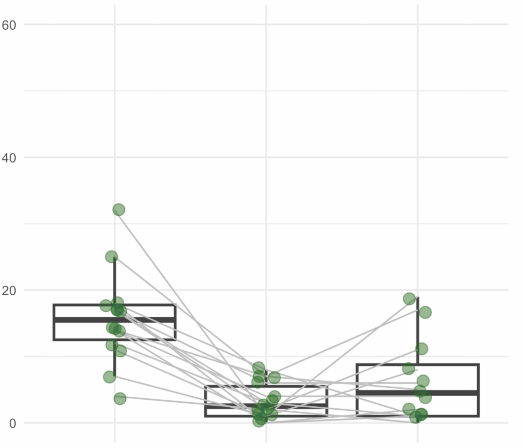

Aggression to Neighbor (per 10min)

**Habituation**

**Stimulus**

**Treatment**

**Habituation**

**Stimulus**

**Treatment**

**D)**

**E)**

**Day 1**

**Day 4**

**Same Neighbor-New Location**

**Day 1**

**Day 4**

**New Neighbor-Same Location**

**Supplementary Figure 3. Male aggression to neighbor from habituation to each treatment.** A) Males from the Same Neighbor-Same Location Treatment. B) Males from the Same Neighbor-New Location Treatment. C) Males from the New Neighbor-Same Location Treatment. D) Males from the Same Neighbor-New Location during cue competition experiments E) Males from New Neighbor-Same Location from cue competition experiments. Gray lines connect the same males from habituation to stimulus treatment and asterisks represent significant differences (*p* < 0.05) based on mixed models.

**New Neighbor-Same Location**

**Day 4**

**Day 1**

**Same Neighbor-New Location**

**Day 4**

**Day 1**

**Day 1**

**Day 4**

**Same Neighbor-Same Location**

*

*

*

*

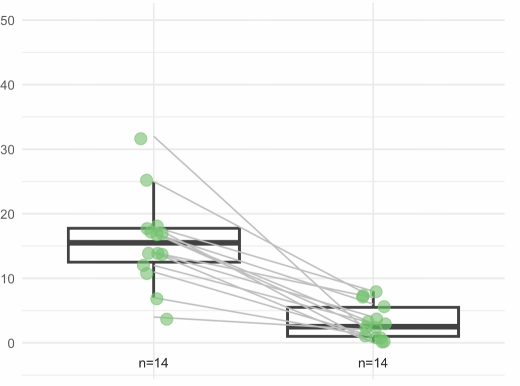

Aggression to Neighbor (per 10min)

**Day 1**

**Day 4**

*

**Supplementary Figure 4: Experiment 3- Neighbor directed aggression during habituation.** Aggression decreased day 1 of habituation compared to day 4 in the simultaneous stimuli treatment. Gray lines connect the same males on day 1 and day 4. Asterisks represent significant differences (*p* < 0.05).
